# The secreted tyrosine kinase VLK is essential for normal platelet activation and thrombus formation

**DOI:** 10.1101/2021.01.24.427281

**Authors:** Leila Revollo, Glenn Merrill-Skoloff, Karen De Ceunynck, James R. Dilks, Mattia Bordoli, Christian G. Peters, Leila Noetzli, Andreia Ionescu, Vicki Rosen, Joseph E. Italiano, Malcolm Whitman, Robert Flaumenhaft

## Abstract

Tyrosine phosphorylation of extracellular proteins is observed in cell cultures and in vivo, but little is known about the functional roles of tyrosine phosphorylation of extracellular proteins. Vertebrate Lonesome Kinase (VLK) is a broadly expressed secretory pathway tyrosine kinase present in platelet ɑ-granules. It is released from platelets upon activation and phosphorylates substrates extracellularly. Its role in platelet function, however, has not been previously studied. In human platelets, we identified phosphorylated tyrosines mapped to luminal or extracellular domains of transmembrane and secreted proteins implicated in the regulation of platelet activation. To determine the role of VLK in extracellular tyrosine phosphorylation and platelet function, we generated mice with a megakaryocyte/platelet-specific deficiency of VLK. Platelets from these mice are normal in abundance and morphology, but have dramatic changes in function both in vitro and in vivo. Resting and thrombin-stimulated VLK-deficient platelets demonstrate a significant decrease of several tyrosine phosphobands. Functional testing of VLK-deficient platelets shows decreased PAR4- and collagen-mediated platelet aggregation, but normal responses to ADP. Dense granule and α-granule release are reduced in these platelets. Furthermore, VLK-deficient platelets exhibit decreased PAR4-mediated Akt (S473) and Erk_1/2_(T202/Y204) phosphorylation, indicating altered proximal signaling. In vivo, mice lacking VLK in megakaryocytes/platelets demonstrate strongly reduced platelet accumulation and fibrin formation following laser-injury of cremaster arterioles compared to controls. These studies demonstrate that the secretory pathway tyrosine kinase VLK is critical for stimulus-dependent platelet activation and thrombus formation, providing the first evidence that a secreted protein kinase is required for normal platelet function.

## Introduction

Phosphorylation of extracellular proteins has been recognized for over a century,^1^ and a large fraction of proteins secreted into the extracellular space are phosphorylated.^2^ Yet only recently have investigators begun to understand the role that phosphorylation of extracellular proteins serves in modulating protein function outside the cell. An important reason for the gap in knowledge regarding the functional significance of phosphorylation of extracellular proteins was the delay in identifying extracellular kinases. Within the last decade, bioinformatic strategies, coupled with biochemical approaches, have led to the identification of kinases with signal sequences that direct them to the secretory pathway and outside the cell, where they can phosphorylate transmembrane and secreted substrates. Such secretory kinases include serine/threonine kinases such as Fam20C^3^ and Drosophila four-jointed (fj),^4^ sugar kinases such as Fam20B^5^ and SGK196,^6^ and the tyrosine kinase VLK/PKDCC, herein referred to as VLK.^7^ Of these secretory kinases, only VLK has been confirmed to reside in platelets.^7^

VLK is a ~54 kD glycoprotein with a 32 amino acid hydrophobic sequence that targets VLK to the secretory pathway and the extracellular environment.^7–9^ As a kinase, it shows a significant preference for tyrosine, however, no consensus sequence for its phosphorylation has been identified.^7^ The catalytic activity of VLK appears to be necessary for its own secretion, since kinase dead mutants are not secreted. It has diverse cellular functions such as mediating axonal branching by phosphorylating repulsive guidance molecule b,^9^ regulating Hedgehog signaling,^10^ modifying phosphorylation of extracellular matrix proteins in the trabecular meshwork of the eye,^11^ and phosphorylating several components of the ER-proteostasis machinery.^12^ Targeted disruption of the gene that encodes VLK (*i.e., Pkdcc*) in mice results in perinatal death and impaired skeletal, intestinal and lung development.^8,13^ Genome wide association studies (GWAS) associate VLK with increased fracture risk in humans^14^ and flight efficiency in birds.^15^ Biallelic gene disrupting variants in VLK have been identified in two individuals with marked skeletal abnormalities.^16^ Although regulated secretion of endogenous VLK was first established in platelets,^7^ and evaluation of the platelet phosphotyrosine proteome reveals phosphorylation of several secreted proteins and extracellular domains of membrane proteins (**Supplementary Table 1**),^17–20^ the effect of VLK deficiency on platelet function has not previously been evaluated.

In platelets, VLK is secreted in an activation dependent manner and is responsible for the phosphorylation of several extracellular proteins.^7^ To understand the contribution of tyrosine phosphorylation of secreted factors and extracellular domains of transmembrane proteins in platelet function, we have examined the effects of loss of VLK in platelets. Mice with a megakaryocyte/platelet-specific deficiency of VLK were generated, and their platelets evaluated. Platelets from these mice were normal in number and morphology, but defective in phosphorylation of tyrosine and stimulation-dependent aggregation. Mice lacking platelet VLK demonstrated normal bleeding times, but a substantial defect in thrombus formation following laser induced damage of cremaster arterioles. These results demonstrate a novel role for the secretory pathway tyrosine kinase VLK in platelet function and thrombus formation, and provide the most compelling evidence to date that secreted kinases contribute to platelet function.

## Material and methods

### LC-MS/MS

Human platelets were isolated from blood collected from 12 healthy donors as described in Supplementary Methods. Cell pellets were solubilized with RIPA buffer followed by acetone (MiliporeSigma) precipitation. Lysates were processed as previously described for phosphotyrosine site identification at the Mass Spectrometry Core in Beth Israel Deaconess Medical Center, Boston MA, USA.^21^ Tryptic digests were injected to an Orbitrap Elite mass spectrometer (ThermoFisher Scientific) using an EASY-NLC II nanoflow HPLC. Raw MS/MS fragmentation data was analyzed with Andromeda integrated to MaxQuant software with a false discovery rate (FDR) of 1% versus the human protein database.

### Cell culture, transfection, and cloning

293T cells (ATCC) were grown in Dulbecco’s modified Eagle’s medium (DMEM) supplemented with 10% (v/v) fetal bovine serum (FBS) and 1X penicillin/streptomycin/amphotericin B (Lonza) at 37°C with 5% CO_2_. Cells were transfected with Lipofectamine 2000 (ThermoFisher Scientific) according to manufacturer protocol. All VLK plasmids and synovial K4 cell lines have been previously described.^7^ Human ENTPD6 cDNA (SinoBiological) lacking the stop codon was subcloned into plasmid modified from pCS2+ expression vector containing a C-terminal V5 tag to generate a plasmid expressing ENTPD6-V5.

### Protein isolation, purification, and immunoblotting

Cells were lysed in RIPA buffer with protease and phosphatase inhibitors. For RIPA-resistant extracts, remaining pellets after RIPA extraction were washed and reconstituted in 1% sodium dodecyl sulfate (SDS). Lysates were immunoprecipitated with anti-V5 beads (Sigma-Aldrich) and eluted with epitope-specific peptide (Tufts University Core). Phospho-tyrosine immunoprecipitations were conducted with anti phosphotyrosine (anti p-Tyr) agarose-coupled beads clone 4G10 (Millipore Sigma) and sepharose-coupled beads P-Tyr-1000 (Cell Signaling Technology). Phenyl-phosphate (PP) (Sigma-Aldrich) was used to control non-specific binding to beads. VLK was immunoprecipitated with antibody raised against GELKVTDLDDARVEETPC epitope of VLK (Abfrontier) conjugated to protein A agarose and eluted with epitope-specific peptide (Tufts University Core). Eluate was deglycosylated with PNGase-F (New England BioLabs) according to manufacturer specifications. Samples were examined by immunoblotting after SDS-PAGE with 8%, 16% or 4-12% gradient gels (ThermoFisher Scientific). The following primary antibodies were used for protein immunoblots: Anti p-Tyr (4G10, Millipore Sigma, #05-321 and P-Tyr-1000, Cell Signaling Technology, #8954), anti V5-HRP (ThermoFisher Scientific, #46-0708), anti ENTPD6 (GeneTex, #GTX101851), anti VLK raised against epitope SRAEYQRIPDSAITQEDYR of mouse VLK (Abfrontier), anti p-VLK (Y64) raised against epitope GRGELARQIRERYEEVQRYSRG phosphorylated at Y64 of mouse VLK (Abfrontier), anti cytoplasmic actin (Milipore Sigma, #A4700), anti GAPDH-HRP (GeneTex, #GTX627408-01), anti p-Akt (S473) (Cell Signaling Technology, #4060), anti Akt (Cell Signaling Technology, #4691), anti p-Erk_1/2_(T202/Y204) (Cell Signaling Technology, #4370), anti Erk_1/2_(Cell Signaling Technology, #4695), anti P-selectin (R&D Systems, #AF737), anti ITGβ3 (Cell Signaling Technology, #13166), anti ITGα2b (Boster, #PB9647), anti FGNγ (GeneTex, #GTX108640), anti vWR (Dako, #A0082 and Santa Cruz Biotechnology, #sc-271409), anti Platelet Factor 4 (PF4) (R&D Systems, #AF595), anti GPIbɑ (Emfret Analytics, #M043-0), and anti thrombospondin 1 (Tsp1) (Bethyl, #A304-989A-M). Image acquisition was done with PXi4 Chemiluminescent and Fluorescent Imaging System (Syngene). Quantification of percent protein phosphorylation was done with Adobe Photoshop 2020.

### Transgenic mice

*Vlk* ^f*lox; neo*^ mice^22^ were a generous gift from Dr. Aimée Zuniga. Introduction of a neo cassette in the reverse orientation in the Vlk locus of *Vlk* ^f*lox; neo*^ mice renders the *Vlk* allele hypomorphic resulting in perinatal lethal homozygous mice. To remove the neo cassette, *Vlk* ^f*lox; neo*^ mice were crossed with FLP1 mice (JAX stock # 003946). FLP1-mediated recombination resulted in deletion of the NEO cassette flanked by *frt*-sequences in the resulting offspring (*FLP1; Vlk ^flox; neo^* mice), which were then mated with wild type mice to remove the FLP1 transgene. The resulting heterozygous progeny *(Vlk ^flox / +^*) were subsequently mated to obtain homozygous *Vlk^flox / flox^* mice herein referred to as *Vlk^f/f^*. *Vlk^f/f^* mice are viable, morphologically normal, born with correct Mendelian frequency, and fertile. To conditionally ablate *Vlk* in cells of the megakaryocytic lineage, *Vlk^f/f^* female mice were bred to male mice expressing *Cre* under the control of the platelet factor 4 (Pf4) promoter (*Pf4-Cre*)^23^ resulting in *Vlk^f/f^*;*Pf4-Cre* mice (*Vlk-cKO*). For genotyping, PCR was performed on genomic DNA isolated from mouse tails by HotSHOT method.^24^ Primers to detect *Vlk^fl^* allele were: 5.4 F, 5’-cacacgctcaatcataccacacc-3’ and 3.7 R, 5’-ggtcattaggtcacagggtaggg-3’. They yield 2 PCR products at 350 bp and 241 bp corresponding to the flox or wildtype allele, respectively. Primers to detect *Pf4-Cre* allele were: F, 5’-cccatacagcacaccttttg-3’ and R, 5’-tgcacagtcagcaggtt-3’.^23^ They yield a single PCR product at 450 bp corresponding to *Pf4-Cre* amplicon. All experiments were performed with 10-14 weeks old male mice, and littermate *Vlk^f/f^* mice were used as controls. Mice were housed in a room maintained at 25°C, on a 12-hour light/dark cycle, and fed regular chow ad libitum. All procedures involving mice were done under the approval of the Harvard Medical Area Institutional Animal Care and Use Committee.

### Intravital microscopy of laser-induced arteriolar thrombosis

Intravital microscopy of microcirculation in the cremaster arteriole was done using laser-injury as described before.^25^ Mice were anesthetized with pentobarbital (5 mg/kg) administered via jugular vein cannula to maintain anesthesia throughout the procedure. Fibrin generation and platelet accumulation were detected at the injury site with Dylight 488-labeled anti-fibrin antibody (59D8, 0.5 μg/g body weight) and Dylight 649-labeled anti-CD42b antibody (M040-3, 0.1 μg/g body weight) injected through jugular vein catheter. Median fluorescence values over time in >30 thrombi in 3-4 mice per genotype were analyzed using Slidebook 6.0 (Intelligent Imaging Innovations).^26–28^ Area under the curve (AUC) was calculated for individual thrombi and compared between *Vlk^f/f^* and *Vlk-cKO* mice to evaluate statistical significance.

Methods for electron microscopy, platelet isolation, platelet aggregation, dense granule secretion assay, P-selectin surface expression assay, tail bleed assay and statistical analysis are provided in Supplementary Methods.

## Results

### Tyrosine phosphorylation of luminal or extracellular domains of transmembrane and secreted proteins in platelets

In order to study phosphorylation on tyrosine residues in extracellular domains of transmembrane and secreted proteins in platelets, we first detected proteins that are tyrosine phosphorylated in platelet lysates in an activation-dependent manner, and subsequently used an informatics-based approach to identify specific proteins and their respective phosphosites. Platelets were stimulated using SFLLRN and phosphoproteins were then immunoprecipitated using an anti-phosphotyrosine (anti-p-Tyr) antibody. Immunoprecipitation performed in the presence of phenyl phosphate (PP), a competitive inhibitor of anti-p-Tyr binding^29^ was used to assess non-specific binding to beads, which was negligible (**Fig. 1A**). Proteins that became phosphorylated in response to PAR1 activation were identified by mass spectroscopy. This analysis identified 213 unique phosphopeptides (**Supplementary Table 2**). Within this pool, 4 tyrosine phosphosites were mapped to proteins with signal peptides or extracellular domains on transmembrane proteins as annotated in UniprotKB (**Fig. 1B**). Among the tyrosine phosphoproteins identified, we focused our attention on ectonucleoside triphosphate diphosphodydrolase 6 (ENTPD6/CD39L2), a member of the CD39/NTPDase family of ectonucleotidases known to regulate platelet activation and thrombus formation.^30,31^ Our study revealed Y290, a residue conserved among NTPDase family members, as a site phosphorylated in ENTPD6 in activated platelets (**Fig. 1B and Supplementary Table 2**). Evaluation of platelet lysates demonstrated that a band corresponding to the molecular weight of ENTPD6 is phosphorylated at tyrosine in an activation dependent manner (**Fig. 1C**). Of note, the total ENTPD6 band in platelet lysates is decreased following stimulation with the PAR1 agonist, SFLLRN, suggesting secretion of the enzyme. Anti-p-Tyr immunoprecipitates from both resting and stimulated platelets identify ENTPD6, confirming that it is tyrosine phosphorylated (**Fig. 1C**). Since VLK is a secretory tyrosine kinase found in platelet α-granules,^7^ we evaluated whether co-transfection of 293T cells with VLK and ENTPD6-V5 increased phosphorylation of the enzyme. Analysis of V5 immunoprecipitates from detergent extracted cells showed increased phospho-tyrosine in the presence of wild-type VLK, but not in the presence of a kinase-dead mutant VLK (**Fig. 1D**).

**Figure 1.**
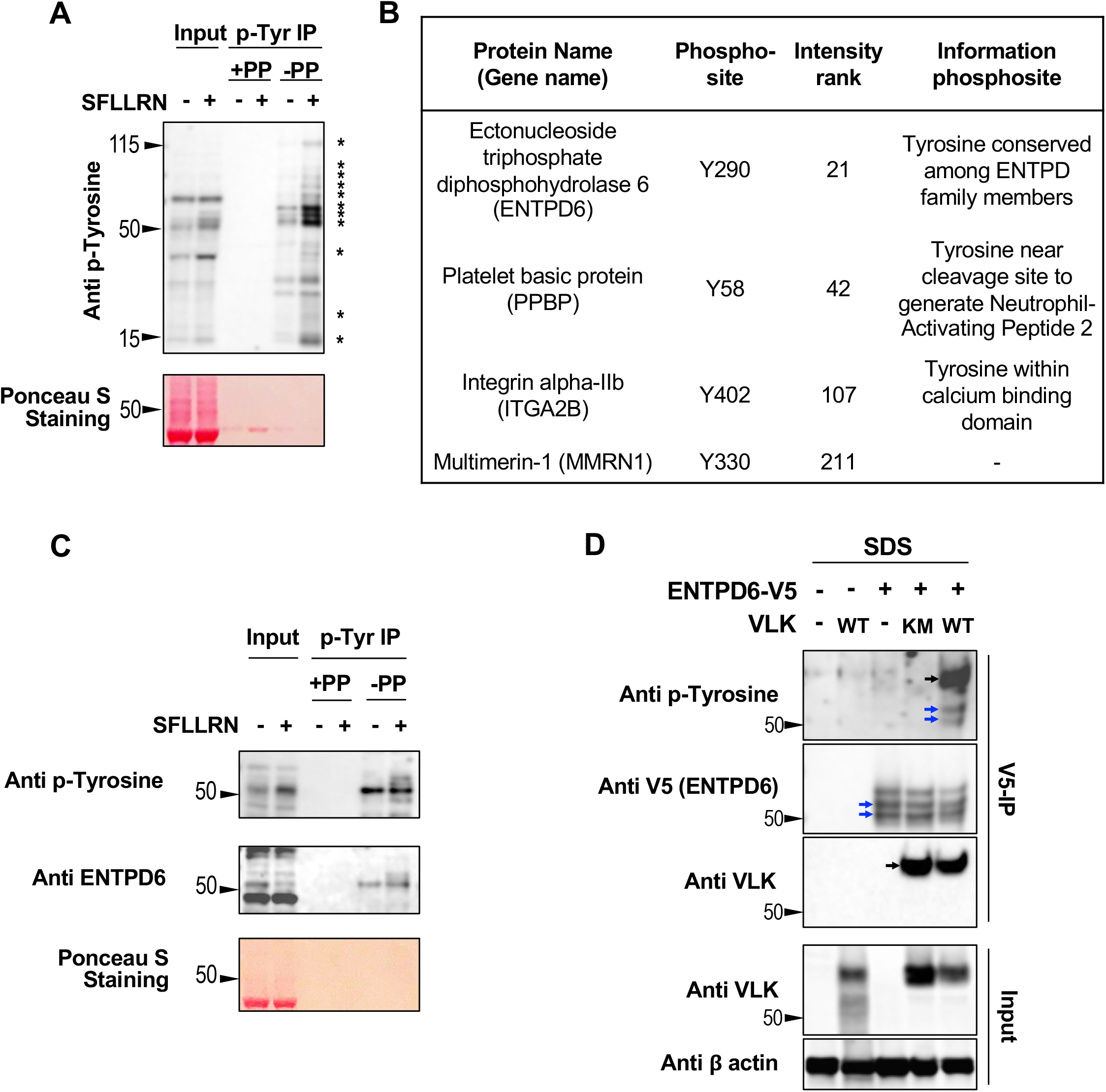
Evaluation of extracellular tyrosine phosphorylation of proteins in human platelets. **(A)** p-Tyrosine (p-Tyr) protein levels in total (Input) and p-Tyr immunoprecipitated fraction (p-Tyr IP) from platelet lysates were assessed by Western blot analysis. Phenylphosphate (PP) was used as negative control for non-specific binding to p-Tyr beads and Ponceau S staining was used to examine total protein levels in input samples. Asterisks indicate activation-dependent tyrosine phosphorylation. **(B)** Phosphorylated tyrosine peptides isolated by anti-p-Tyr IP of tryptic digests from TRAP-stimulated human platelet lysates were analyzed by LC-MS/MS. A total of 213 unique phosphopeptides were identified; within this pool, 4 tyrosine phosphosites were mapped to proteins with signal peptides or extracellular domains on transmembrane proteins as annotated in UniprotKB. (**C**) ENTPD6 (CD39L2) levels were determined in total (Input) and p-Tyr IP fraction in lysates from human platelets. Total protein levels in input were determined with Ponceau S staining. (**D**) Tyrosine phosphorylation of V5 immunoprecipitates in RIPA-resistant extracts (SDS) from 293T cells co-expressing V5-tagged ENTPD6 with wildtype (WT) or kinase dead (KM) VLK. Extracts (Input) were analyzed for VLK and actin. Blue and black arrows indicate bands corresponding to ENTPD6 and VLK, respectively.

### Platelet/Megakaryocyte-specific deletion of VLK in mice

VLK deficient mice die within a day after birth.^9,13^ In order to evaluate the function of platelets that lack VLK, we crossed VLK floxed mice (*Vlk^f/f^*) with mice expressing the Cre recombinase under the control of the PF4 promoter (PF4-Cre mice) resulting in *Vlk^f/f^*;*Pf4-Cre* mice (*Vlk-cKO*). Analysis of genomic DNA from *Vlk-cKO* mice demonstrated the presence of the targeted loxP and Pf4-Cre alleles (**Fig. 2A**). *Vlk-cKO* mice were born in normal numbers and survived until adulthood. They were normal weight and did not have gross morphological defects. Platelets isolated from these mice demonstrated a lack of VLK as evaluated by Western blot analysis (**Fig. 2B**). In contrast, P-selectin and GAPDH levels were indistinguishable between control and VLK knockout platelets, indicating equal protein loading. Complete blood counts (CBC) were performed to identify any gross abnormalities in hematopoiesis resulting from platelet-specific deficiency of VLK. No significant differences in leukocyte, erythrocyte, or platelet counts were observed between *Vlk-cKO* mice and controls (**Supplementary Table 3**). Similarly, no differences in mean platelet volume were observed. Evaluation of platelets by transmission electron microscopy demonstrated normal platelet morphology and did not indicate any obvious differences in platelet shape or organelle number (**Fig. 2C**). Dense granules and α-granules in VLK-deficient platelets were intact and were similar in size and morphology to controls. Western blot analysis of lysates from wild-type and VLK-deficient platelets demonstrated a normal complement of the major platelet surface receptors α_2b_β_3_, GPIbα, and P-selectin (**Fig. 2D**) and the major secretory proteins fibrinogen, von Willebrand factor, and platelet factor 4 (PF4) (**Fig. 2E**).

**Figure 2.**
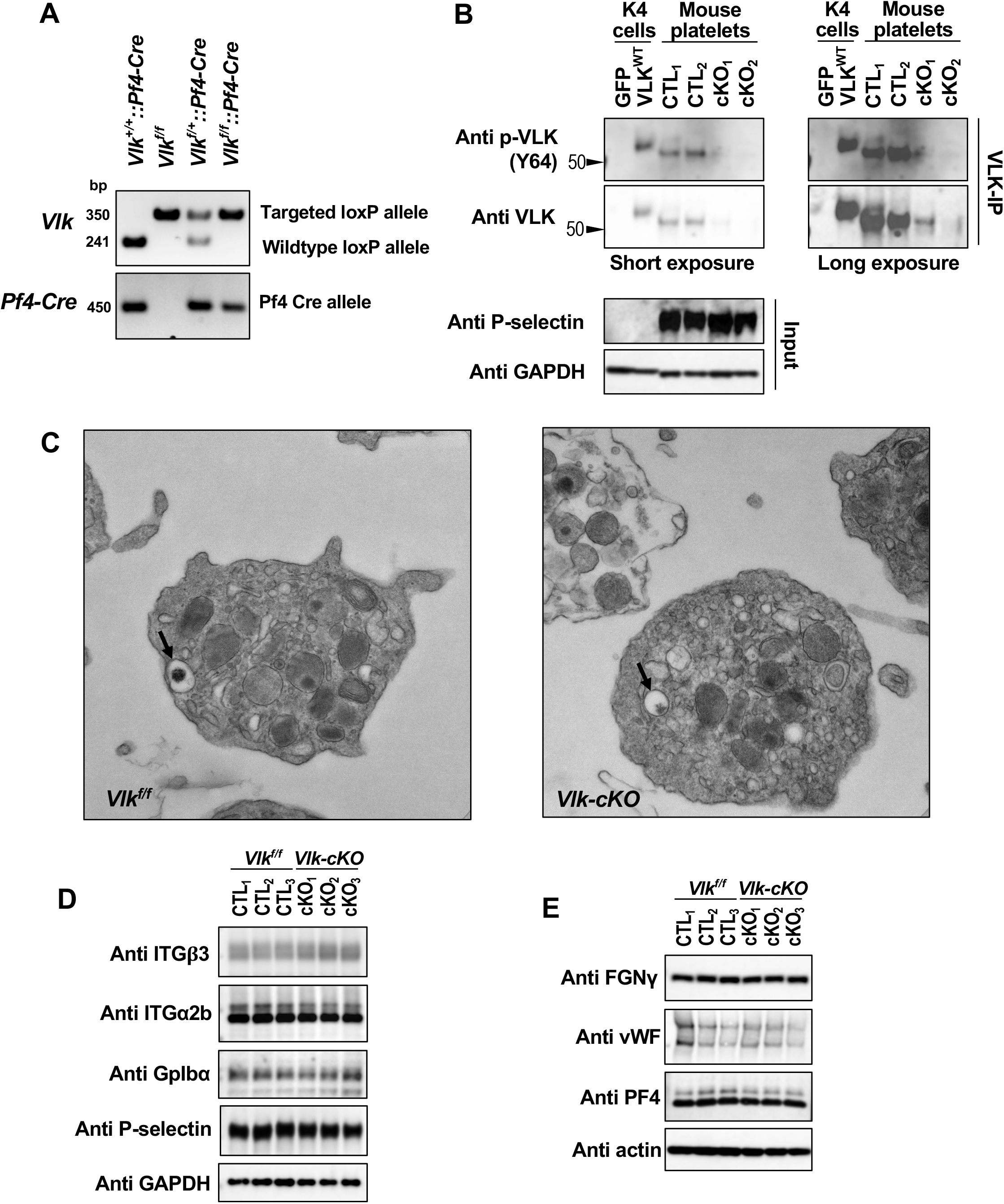
*Vlk* deficiency in platelets does not affect platelet morphology or levels of platelet receptors and cargo proteins. **(A)** Analysis of genomic DNA from *Vlk^+/+^::Pf4-Cre, Vlk^f/f^, Vlk^f/+^::Pf4-Cre,* and *Vlk^f/f^::Pf4-Cre* mice. Targeted *loxP* allele (*fl*) is 350 bp; *wildtype* allele (*+*) is 241 bp; *Pf4-Cre* allele is 450 bp. **(B)** Western blot analysis of phosphorylated and total VLK immunoprecipitates (VLK-IP) prepared from *Vlk^f/f^* (CTL) and *Vlk-cKO* (cKO) washed platelet lysates. Lysates from K4 synoviocytes stably expressing GFP or wildtype mouse VLK (VLK^WT^) were used as controls for VLK protein. Anti P-selectin and GAPDH were used as controls for platelet-specific and total protein levels in input samples, respectively. **(C)** Representative image of electron microscopy analysis of resting *Vlk^f/f^* and *Vlk-cKO* platelets. Dense granules indicated by black arrows. Representative images were obtained at 13,000x. **(D, E)** Protein levels in platelets from *Vlk^f/f^* (CTL) and *Vlk-cKO* (cKO) mice (n=3 per genotype). Anti-GAPDH and actin antibodies were used to examine total protein levels. **(D)** Platelet receptors integrin β_3_, integrin α_2b_, glycoprotein Ibα, and P-selectin. **(E)** Platelet cargo fibrinogen, von Willebrand factor, and platelet factor 4.

To determine whether deficiency of VLK in platelets affects activation-dependent tyrosine phosphorylation, control and VLK-deficient platelets were stimulated with thrombin to elicit full activation. Secretory proteins including Tsp1 and PF4 were evaluated in the lysates and supernatants to demonstrate that VLK-deficiency did not alter release of granule contents under these conditions of full activation (**Fig. 3A**). Phosphoproteins phosphorylated at tyrosine residues were then immunoprecipitated from lysates of control and VLK-deficient platelets before and after thrombin stimulation. Western blot analysis demonstrated several proteins that were tyrosine phosphorylated in resting platelets (**Fig. 3B,*a-f***) and following platelet activation (**Fig. 3B,*1-10***). Specificity of the immunoprecipitation was demonstrated by inhibition using phenyl phosphate. Comparison of band intensities from several representative bands obtained from Western blots of resting platelets showed decreased phosphorylation of numerous proteins in VLK-deficient platelets relative to controls (**Fig. 3C**). Analysis of tyrosine phosphorylation of activated platelets also showed decreased phosphorylation in VLK-deficient platelets compared to control (**Fig. 3D**). The secreted protein Tsp1 has been previously identified as a tyrosine phosphorylated protein in platelets (**Table S1**). A significant decrease in tyrosine phosphorylation of Tsp1 was observed in VLK-deficient platelets when compared with controls (**Figs. 3E, F**). These studies demonstrate that tyrosine phosphorylation is decreased in platelets deficient in VLK and the changes in tyrosine phosphorylation in *VLK-cKO* platelets include secreted proteins such as Tsp1.

**Figure 3.**
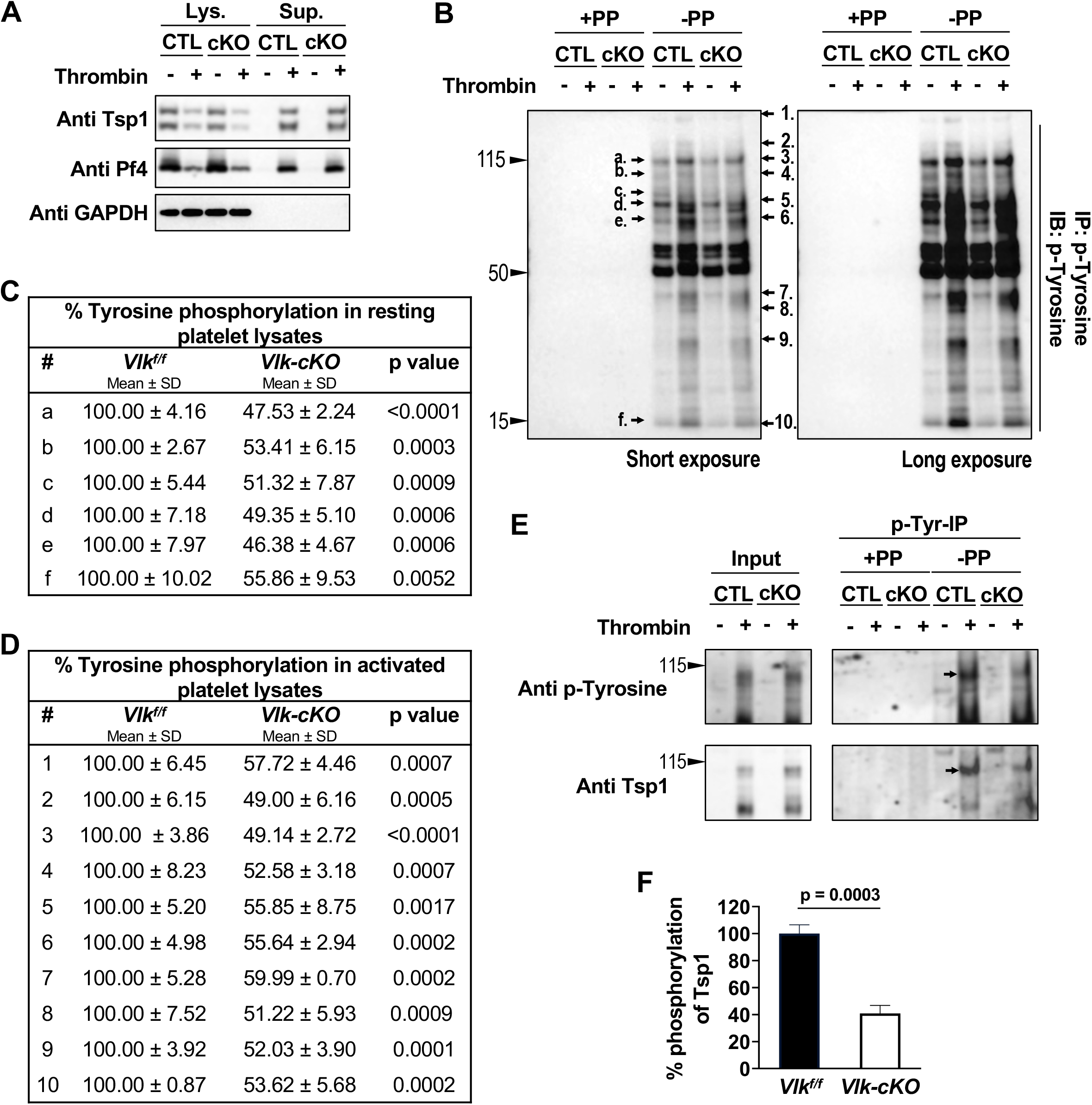
Reduced tyrosine phosphorylation in resting and activated mouse platelets harboring deletion of *Vlk* compared to control. **(A)** Immunoblots of lysates (Lys.) and supernatants (Sup.) from *Vlk^f/f^* (CTL) and *Vlk-cKO* (cKO) platelets resting or stimulated with 5 U/mL thrombin for 15 minutes. Anti-Tsp1 and anti-PF4 antibodies were used as markers of platelet activation, and anti-GAPDH antibody was used to examine total protein levels in lysates. **(B)** Lysates from A were used to examine tyrosine phosphoproteins by immunoblot (IB) of p-Tyr enriched fraction (IP). Black arrows indicate phopshobands decreased in *Vlk-cKO* (cKO) compared to *Vlk^f/f^* (CTL) in resting (a-f) and activated (1-10) platelets. **(C, D)** Cumulative percent tyrosine phosphorylation in phosphobands indicated in representative figure in **B** from resting (**C**) and activated (**D**) control and *Vlk-cKO* lysates; data are expressed as mean ± SD; n=3 independent experiments. **(E)** Representative image of input and p-Tyr IP fraction from resting and thrombin-stimulated supernatants analyzed by SDS-PAGE with 8% gels and immunoblotting with anti-p-Tyr and anti-Tsp1 antibodies. Phenylphosphate (PP) was used as negative control for non-specific binding to p-Tyr beads. Black arrows indicate bands decreasing in cKO compared to CTL. **(F)** Cumulative percent phosphorylation of Tsp1 in activated supernatants from *Vlk-cKO* compared to CTL; n=3 independent experiments.

### Role of VLK in platelet function

The observation that VLK-deficient platelets showed normal secretion of α-granule cargos in response to thrombin raises the possibility that platelet function is unaffected by the absence of VLK. To evaluate this possibility more rigorously, we compared aggregation of VLK-deficient platelets to control platelets using a variety of agonists. In contrast to the full activation observed in response to thrombin, VLK-deficient platelets showed reduced aggregation in response to stimulation of PAR4 with 100 μM AYPGKF compared to *Vlk^f/f^* controls (*Vlk^f/f^*: 70±5.1% aggregation [n=10]; *Vlk-cKO*: 23±8.0% aggregation [n=10]) (**Fig. 4A**). Aggregation in response to stimulation with 4 μg/mL collagen was also reduced in VLK-deficient platelets (*Vlk^f/f^*: 53±2.5% aggregation [n=4]; *Vlk-cKO*: 27.5±2.9% aggregation [n=4]) (**Fig. 4B**). The defect in platelet aggregation in VLK-deficient platelets was overcome in response to 5 U/ml thrombin (**Supplementary Fig. S1**). A dose curve of platelet α-granule release as detected by PAR4-mediated appearance of P-selectin at the surface demonstrated impaired responses at 100 μM AYPGKF, but was overcome at 150 μM AYPGKF (**Fig. 4C**). Release of α-granule cargo was also decreased in *Vlk-cKO* platelets, as evidenced by reduction of Tsp1 and PF4 secretion in response to 100 μM AYPGKF (**Fig. 4D**). Dense granule release in control platelets occurred in response to 50 μM AYPGKF, but was absent in platelets that lack VLK (**Fig. 4E**). Addition of a subthreshold concentration of ADP (2.5 μM) restored platelet aggregation in *Vlk-cKO* platelets (**Fig. 4F**). Furthermore, platelet aggregation in response to ADP was comparable between control and *Vlk-cKO* platelets, regardless of the dose that was used (**Supplementary Fig. S2**). These studies indicate that deficiency of VLK impairs platelet function in response to low to moderate doses of agonists other than ADP.

**Figure 4.**
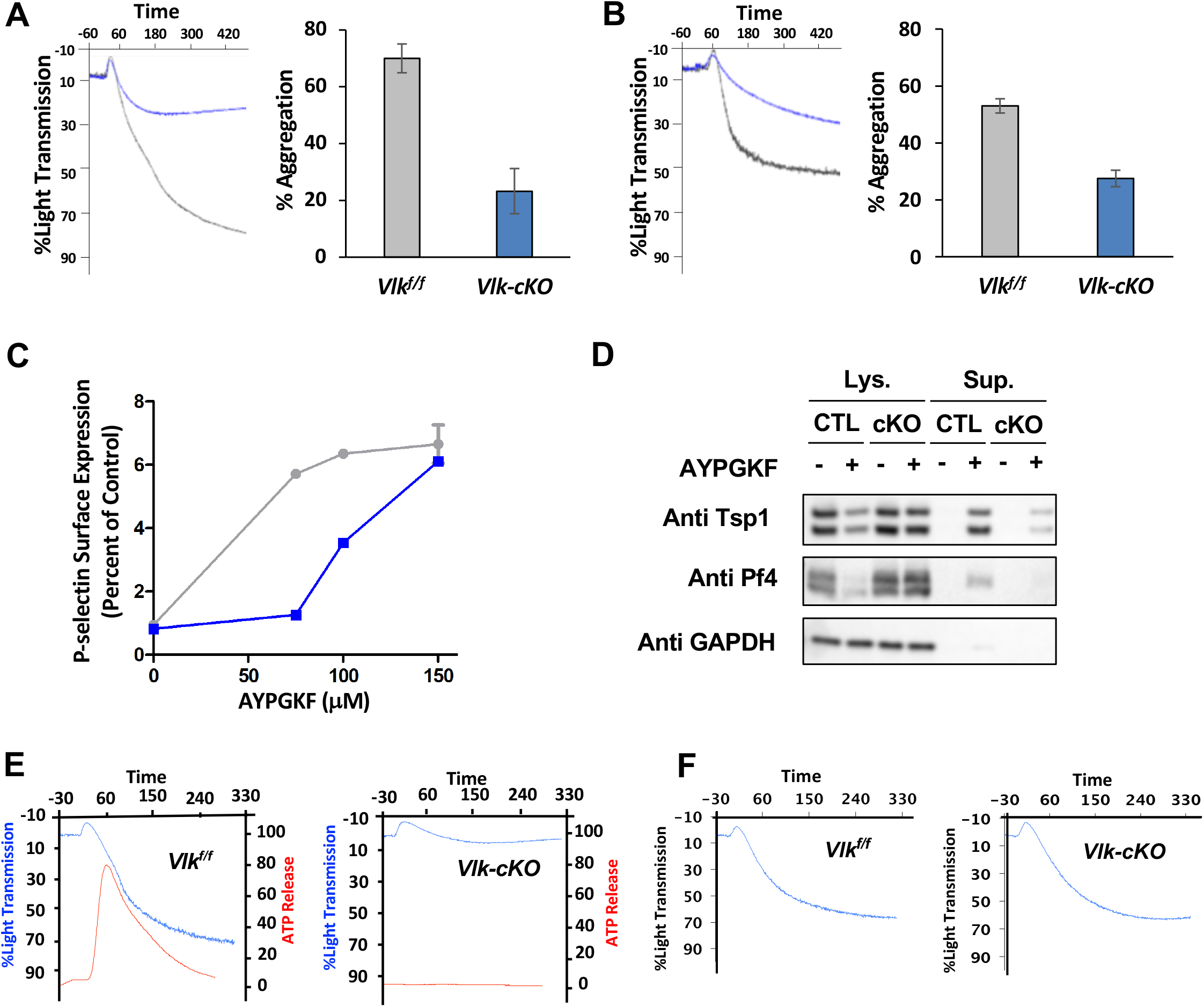
*Vlk* deficiency in platelets alters platelet aggregation and granule exocytosis. **(A)** Evaluation of aggregation of washed platelets (2×10^8^ platelets/ml) from control mice (*grey*) and platelet-specific VLK knockout mice (*blue*) stimulated with 100 μM PAR4 peptide AYPGKF. Representative tracings are shown in *left panel* and cumulative data (n = 3-5) in *right panel*. (**B**) Platelets from control mice (*grey*) and VLK knockout mice (*blue*) were stimulated with 4 μg/mL collagen and evaluated as in **A**. **(C)** Expression of P-selectin in platelets from control (*grey*) and platelet-specific VLK knockout mice (*blue*) in response to AYPGKF. **(D)** Protein immunoblots of lysates (Lys.) and supernatants (Sup.) from *Vlk^f/f^* (CTL) and *Vlk-cKO* (cKO) platelets resting or stimulated with 100 μM PAR4 peptide AYPGKF for 15 minutes. Anti-Tsp1 and anti-PF4 antibodies were used to detect secretion of α-granule cargo, and anti-GAPDH antibody was used to determine total protein levels in lysates. (**E**) Concurrent monitoring of aggregation (*blue trac*ing) and ATP release (*red tracing*) following exposure to 50 μM AYPGKF in platelets from control (*left panel*) and platelet-specific VLK knockout mice (*right panel*). **(F)** Aggregation studies in platelets from control (*left panel*) and platelet-specific VLK knockout mice (*right panel*) were performed as described in **E** except that ADP 2.5 μM was added together with AYPGKF.

To further evaluate the platelet function defect in the absence of VLK, proximal signaling events in response to agonist stimulation were monitored. Exposure of platelets to 100 μM AYPGKF resulted in phosphorylation of Akt (S473) and Erk_1/2_(T202/Y204) (**Fig. 5**). Specifically, Akt (S473) phosphorylation was reduced by 63±18.6% in *Vlk-cKO* compared to control platelets (**Fig. 5A, B**). Erk_1/2_(T202/Y204) phosphorylation was also reduced by 67.6±21.9% in *Vlk-cKO* compared to controls (**Fig. 5C, D**). Taken together, our analysis of platelet function in *Vlk-cKO* platelets indicates that VLK influences release of platelet dense granules and that its absence results in platelets that are less sensitive to platelet activation, but capable of full activation in response to strong agonists (**Fig. 5E**).

**Figure 5.**
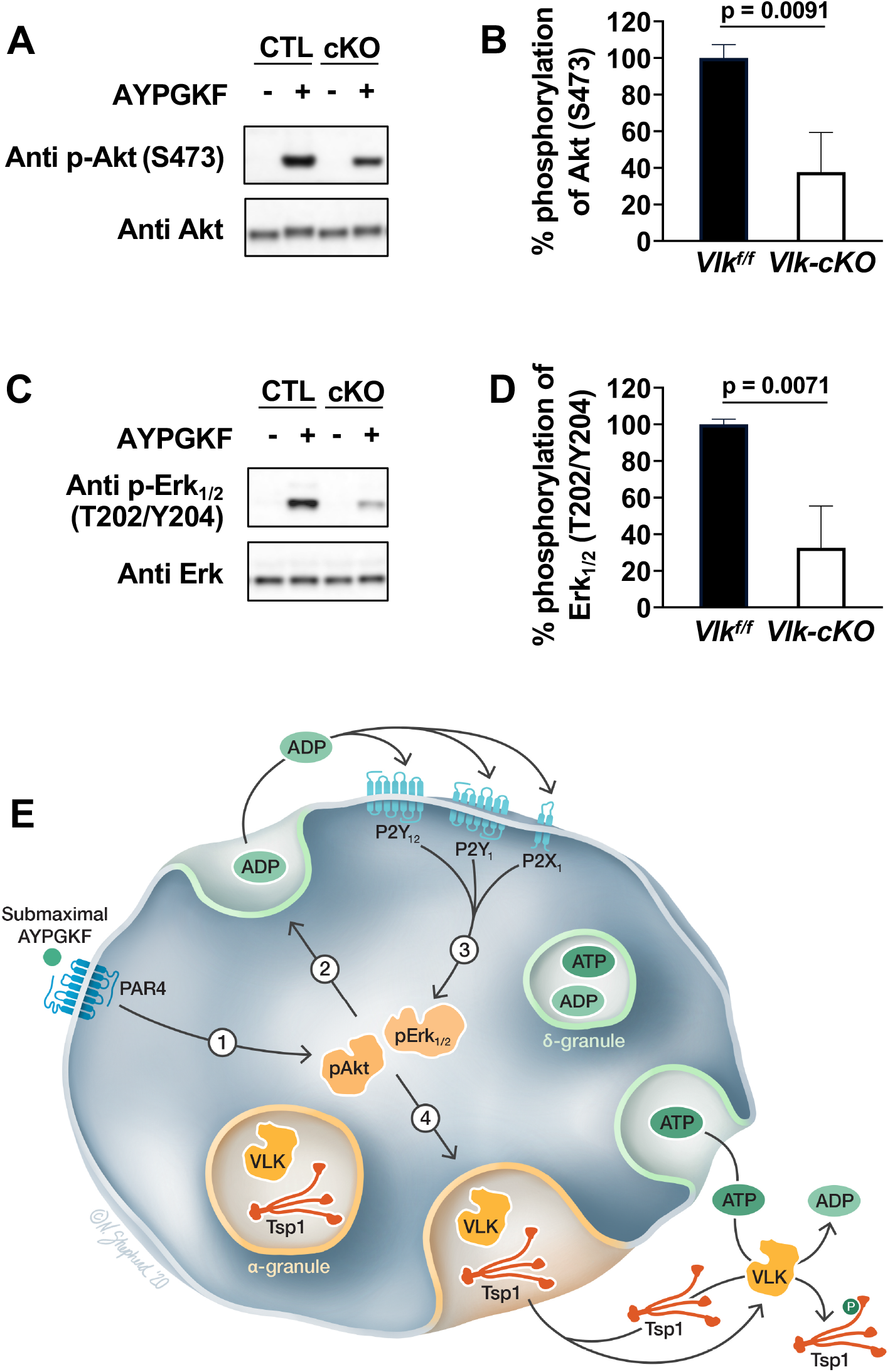
Evaluation of proximal signaling in VLK-deficient platelets. **(A)** Representative image of p-Akt (S473) and total Akt levels in lysates from CTL and cKO platelets resting or stimulated with 100 μM AYPGKF. **(B)** Cumulative percent phosphorylation of S473 in Akt; n=3 independent experiments. **(C)** Representative image of p-Erk_1/2_(T202/Y204) and Erk_1/2_total levels in lysates from CTL and cKO platelets resting or stimulated with 100 μM AYPGKF. **(D)** Cumulative percent phosphorylation of Erk_1/2_; n=3 independent experiments. (**E**) Proposed model for the role of VLK in platelet function. Step 1, stimulation through PAR4 by a submaximal agonist concentration results in partial activation of kinases involved in dense granule release, such as Akt and Erk_1/2_. Step 2, signaling downstream of Akt and Erk_1/2_results in release of ADP from dense granules. VLK appears to act in these steps leading to dense granule release by mechanisms that are yet to be determined but could include phosphorylation of receptors on extracellular domains, phosphorylation of secretory chaperones, or phosphorylation of dense granule resident proteins important for release. Step 3, ADP released from dense granules stimulates purinergic receptors, enhancing signaling through kinases involved in granule release. Step 4, exocytosis of α-granules results in the release of VLK into the extracellular environment along with ATP and VLK substrates, such as thrombospondin 1.

### Platelet VLK functions in thrombus formation *in vivo*

The observation that platelet VLK functions in aggregation and granule release in response to submaximal concentrations of agonists raises the question of whether it is important in thrombus formation. To address this possibility, we evaluated platelet accumulation and fibrin formation following laser-induced injury of cremaster arterioles using intravital microscopy. Platelet accumulation in *Vlk-cKO* mice was reduced to 90% of that of control mice (p ≤ 0.02) (**Figs. 6A, B, D**). In addition, fibrin formation in *Vlk-cKO* mice was reduced to 62% of control mice (p ≤ 0.009) (**Figs. 6A, C, D**). We next evaluated the effect of megakaryocyte/platelet-specific VLK deficiency on tail bleeding times. In contrast to their defective thrombus formation following arteriolar injury, *Vlk-cKO* mice did not demonstrate prolonged tail bleeding times (**Fig. 6E**). Thus, for the assays used in this study, *Vlk-cKO* mice showed a defect in thrombus formation but not bleeding.

**Figure 6.**
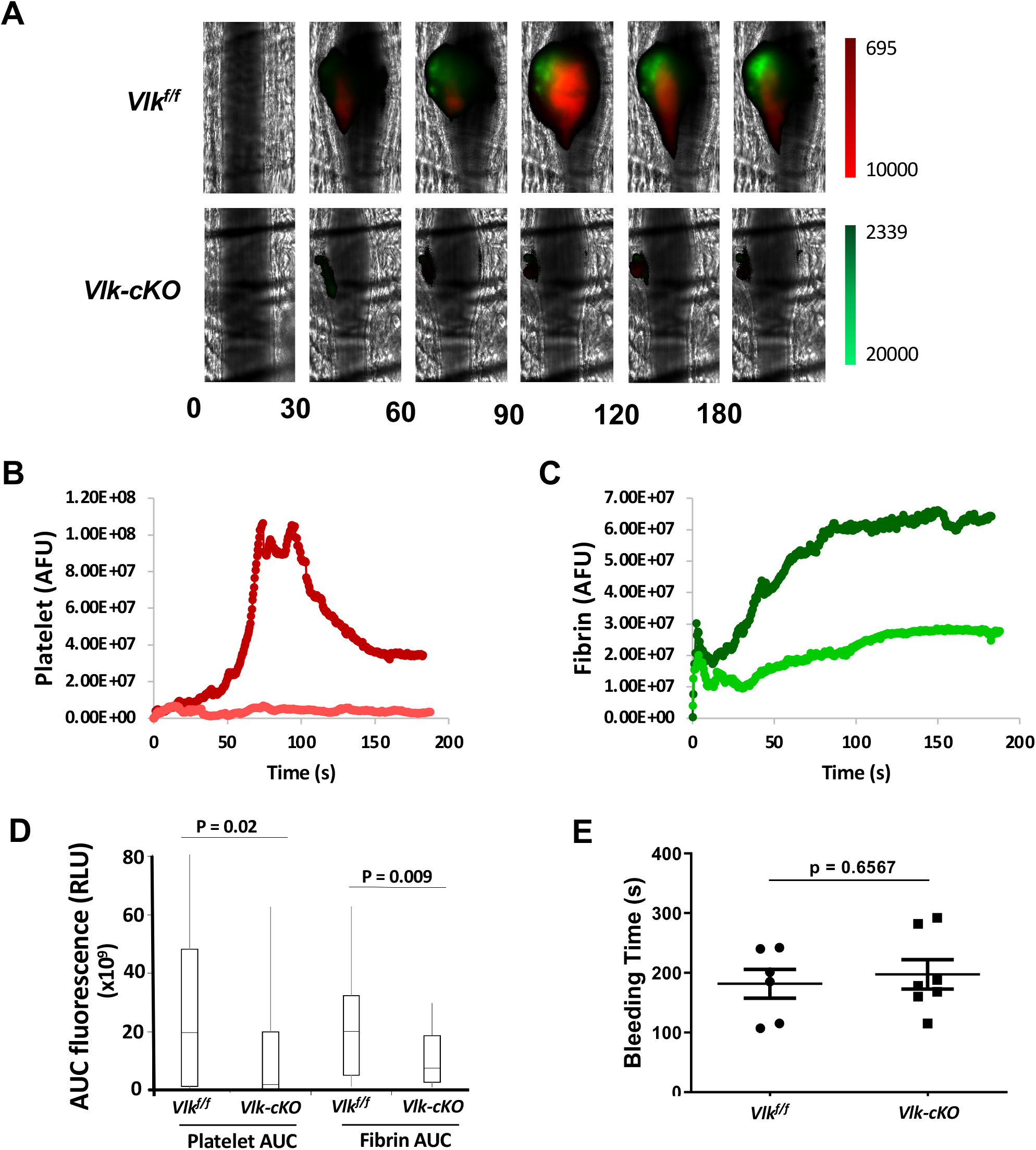
Ablation of *Vlk* in platelet lineage reduces arterial thrombosis without affecting bleeding times. **(A-D)** Cremaster arteriole injury was induced by laser injury in *Vlk-cKO* mice and *Vlk^f/f^* littermate control mice. Platelet and fibrin accumulation at the site of injury were detected for 180 seconds with Dylight 647-labeled platelet-specific anti-CD42b antibody (0.1 μg/g body weight) and Dylight 488-labeled anti-fibrin antibody (59D8, 0.5 μg/g body weight), respectively. **(A)** Representative binarized images from a single thrombus evaluated for the appearance of fluorescence signals associated with platelet accumulation (*red*) and fibrin deposition (*green*) at indicated time points post-injury. **(B, C)** Median integrated **(B)** platelet (*red*) and **(C)** fibrin (*green*) fluorescent intensities following laser injury were calculated for all thrombi in *Vlk^f/f^* (n = 3; 39 thrombi, *darker color*), and *Vlk-cKO* (n = 3; 52 thrombi, *lighter color*) mice. **(D)** Area under the curve (AUC) for (A) platelet and (B) fibrin fluorescent intensities was calculated for all individual thrombi. **(E)** Time to bleeding cessation (seconds) in *Vlk^f/f^* (n = 6), and *Vlk-cKO* (n = 7) mice (unpaired t test, p = 0.6567).

## Discussion

While numerous instances of protein kinase localization and activity in the platelet extracellular environment have been described over the years,^32–35^ the reported kinases (PKA, PKC) are cytoplasmic kinases with no known mechanism for regulated release, and their extracellular localization may be secondary to non-specific cell breakage. Since there have been no genetic approaches to test the role of these putative extracellular activities, their functional significance has remained obscure. We report here the first functional data on a protein kinase that is specifically localized to the secretory pathway in platelets, VLK. We find a role for platelet VLK in aggregation, granule exocytosis, and thrombus formation. Depletion of VLK did not affect platelet count or volume, nor did it cause any gross abnormalities in platelet morphology or changes in blood cell indices (**Fig. 2 and Supplementary Table S3**). Transmission electron microscopy has shown that platelet VLK is sequestered in platelet α-granules.^7^ This localization is consistent with the presence of a N-terminal hydrophobic sequence in VLK that targets it to the secretory pathway. How VLK might be enriched in α-granules will be an interesting area for future investigation. Platelets lacking VLK show no defect in α-granule morphology, number, or secretory pathway protein content, suggesting that VLK is not necessary for normal α-granule biogenesis.

Several lines of evidence point to a dose sensitive defect in stimulus dependent dense granule release in VLK-deficient platelets. Platelets showed impaired aggregation in response to activation by a PAR4 agonist and by collagen, but not in response to ADP (**Fig. 4 and Supplementary Fig. S2**). ADP release from dense granules contributes to stimulation of platelets with submaximal concentrations of PAR4 agonist and collagen, but is not important for ADP-mediated platelet aggregation. Tracings from VLK-deficient platelets showed a lack of dense granule release in response to submaximal concentrations of PAR4 agonist AYPGKF (**Fig. 4E**) and addition of a subthreshold amount of ADP completely reversed the aggregation defect (**Fig. 4F**). Taken together, these observations strongly indicate that dense granule release is impaired in VLK-deficient platelets. A primary defect in dense granule release could also impair α-granule exocytosis.^36^ However, it is not clear that the defect in dense granule release is due to a primary defect in dense granule release, or due to a defect in upstream signaling that is required for granule release. The facts that the initial wave of platelet aggregation (prior to dense granule release) (**Fig. 4E**) and that Akt (**Fig. 5A, B**) and Erk_1/2_(**Fig. 5C, D**) phosphorylation are inhibited in VLK-deficient platelets suggests that the primary defect is upstream of dense granule release.

Our in vivo studies establish that thrombus formation is impaired in *Vlk-cKO* mice following arteriolar injury (**Fig. 6**), consistent with the defects in platelet exocytosis and aggregation we find in purified platelets. It is possible that the observed decrease in fibrin formation (**Fig. 6C, D**) may be secondary to the defect in platelet accumulation (**Fig. 6B, D**), resulting in a decreased surface for fibrin formation. Previous studies using the laser-induced injury model in PAR4-deficient mice have shown, however, that impairment of platelet accumulation does not prevent fibrin formation,^37,38^ suggesting that a platelet aggregation defect may not be sufficient to account for defects in fibrin formation associated with VLK loss. Data pointing to the endothelial monolayer as the primary surface for fibrin formation following laser injury also suggest that the fibrin and platelet aggregation defects result from distinct effects of VLK loss.^38,39^ Interestingly, fibrinogen has been reported to be tyrosine phosphorylated in vivo (**Supplementary Table 1**),^17^ raising the possibility of a more direct effect of VLK on fibrin formation.

Previous work has established that VLK can phosphorylate secreted proteins both in the secretory pathway and following its release from α-granules.^7^ The role of VLK in extracellular phosphorylation following release is particularly intriguing in platelets, because ATP is released from dense granules concomitant with VLK release from α-granules (**Fig. 5E**). We have observed reduced phosphorylation of extracellular tyrosine phosphorylation in Hermansky-Pudlak syndrome platelets, which are deficient in ADP/ATP, suggesting that release of ATP from dense granules is indeed critical for VLK activity in the extracellular environment (**supplementary Fig. S3**). Our findings that VLK-deficient platelets are defective in dense granule release, however, indicate that a key functional defect in these platelets is prior to the stimulus dependent release of ATP, either in the early steps in receptor activation or in stimulus dependent dense granule mobilization. It therefore seems likely that VLK is required in the secretory pathway, either during platelet development or in the resting mature platelet, for normal coupling of agonist stimulation to dense granule release. While VLK depletion results in impaired agonist-stimulated dense granule release (**Fig. 4E**), ADP-mediated activation of purinergic receptors remains unaffected (**supplementary Fig. S2**), indicating that this defect is not due to a generalized defect in platelet activation. Phosphorylation of receptors on extracellular domains, secretory chaperones necessary for receptor surface expression, dense granule resident proteins important for release, are each a possibility for how VLK might be required for normal stimulus-response coupling in platelets. In either case, kinase activity of VLK following stimulus dependent co-secretion with ATP is unlikely to explain this phenotype.

There may be, however, an important role for post release activity of platelet VLK in thrombus formation in vivo. Tyrosine phosphorylation of a broad range of secreted proteins implicated in thrombosis, angiogenesis, and wound healing has been identified in human and mouse tissue samples,^40^ and we have found many of these same phosphorylations both in human platelets (**Fig. 1B**) or following expression of VLK in cultured cells.^7^ Platelets are not the only potential source of VLK in the thrombosis/wound microenvironment, as VLK is present at significant levels in circulating plasma,^41^ and sorting out the full complexity of VLK sources and targets under in vivo circumstances will be a long-term undertaking. Although mice lacking VLK in specific tissues are an essential tool for defining the function of VLK in hemostasis and wound healing, the very small amounts of material available from wound sites in mice are not sufficient for the identification of tyrosine phosphorylated secreted proteins by LC-MS/MS analysis. This limitation currently precludes us for directly identifying candidate VLK target phosphoproteins from sites of local arteriole damage. Further study of tyrosine phosphoproteins and VLK dependent phosphorylations in alternate systems (tissue culture, large animal thromboses) are likely to identify candidate targets that can be interrogated in the murine system.

We have shown that megakaryocyte/platelet-specific deletion of VLK results in platelet function defects and reduced thrombus formation. These observations, particularly the finding that deletion of VLK inhibits thrombus formation without prolonging bleeding times, raise the possibility that pharmacologic inhibition of VLK could represent a novel therapeutic strategy in thrombosis. VLK is widely divergent from cytoplasmic kinases, has extracytoplasmic localization, and post-natal deletion of VLK is well-tolerated (unpublished data), allowing VLK to be targeted distinctively from the rest of the kinome. Detailed phenotyping of resting VLK-deficient platelets demonstrated that the only detectable defect resulting from VLK deficiency is altered tyrosine phosphorylation. Overall, these results demonstrate that VLK is essential for normal platelet function and thrombus formation, and implicate tyrosine phosphorylation of extracellular proteins functions in thrombosis.

## Acknowledgements

The authors thank Nicole Sheperd for creating the illustration in Fig. 5E and Kristen Powers for her assistance generating the ENTPD6-V5 plasmid.

Dr. Whitman has received support from the National Institute of General Medical Sciences (R01GM115417), the National Institute of Arthritis and Musculoskeletal and Skin Diseases (R01AR066717), and National Institute of Dental and Craniofacial Research (R21DE024312). Dr. Flaumenhaft has received support from the National Heart, Lung and Blood Institute (R35HL135775 and R01HL125275). Dr. Italiano has received support from the National Heart, Lung and Blood Institute (R01HL068130 and R01HL136394).

## Authorship

LR designed, performed, and analyzed experiments as well as wrote and edited the manuscript; GMS, KD, JRD, MB, CGP, AI and LN performed experiments and analyzed data; JEI, VR designed experiments and edited the manuscript; MW, RF designed and analyzed experiments, and wrote and edited the manuscript.

