## supplemental materials for "The secreted tyrosine kinase VLK is essential for normal platelet activation and thrombus formation"

### Supplementary Methods

#### Immunofluorescence

Human platelets were isolated from whole blood obtained by venipuncture from a control individual and a patient with Hermansky-Pudlak syndrome (HPS) subtype 1 as previously described,[1] with approval from the Institutional Review Board at Beth Israel Deaconess Medical Center (BIDMC). Cells were stimulated with 150 µM thrombin-receptor activating peptide (TRAP) SFLLRN (Sigma-Aldrich) for 15 minutes or incubated on collagen coated coverslips for 30 minutes. Cells were fixed with 4% paraformaldehyde (PFA) for 15 minutes, blocked overnight with 10% normal goat serum and 1% BSA, incubated with anti- p-Tyr P100 antibody (Cell Signaling Technologies, #9411), and AlexaFluor-488–conjugated secondary antibody (ThermoFisher Scientific). Filamentous actin (F-actin) was stained last using Alexa Fluor 568 Phalloidin (Life Technologies) diluted in methanol to permeabilize and visualize platelets. Coverslips were mounted with Aqua-Poly/Mount (Polysciences) and evaluated by fluorescence microscopy.

#### Electron microscopy

Electron microscopy of mouse platelets was performed as previously described (Battinelli et al., *Blood* 2019). Samples were examined with a Tecnai G2 Spirit BioTWIN electron microscope (FEI company) at an accelerating voltage of 80 kV. Images were recorded with an AMT 2K CCD camera with AMT digital acquisition and analysis software (Advanced Microscopy Techniques). Representative images are shown from platelets isolated from 3 mice per genotype.

#### Platelet isolation

Human and mouse washed platelets were isolated from citrated whole blood by differential centrifugation with HEPES Tyrode buffer in the presence of prostaglandin E1 (PGE-1) as previously described.[2,3] After the last wash, platelets were treated with indicated agonists for 15 minutes at 37ºC, separated from supernatants by centrifugation, and lysed with RIPA buffer (150 mM NaCl, 1% NP40, 0.5% Na-deoxycholate, 0.1% SDS, 50 mM Tris pH 7.5) containing protease and phosphatase inhibitors. For each experimental replicate with mouse platelets, washed platelets were isolated from citrated blood collected by retro-orbital puncture with glass capillaries pooled together from 3 to 5 mice per genotype to ultimately achieve 5x$10^8$ cells per mL.

**Platelet aggregation and dense granule secretion assay**

Platelet aggregation and ATP release from dense granules were measured from washed platelets as described before.[2] Aggregation in response to PAR4-agonist AYPGKF (Sigma-Aldrich), collagen (Chrono-log Corporation), and ADP (Sigma-Aldrich) was determined with ChronoLog 680 aggregometer (Chrono-log Corporation) at indicated concentrations. An agonist dose curve was run to determine the lowest dose at which aggregation was achieved for each independent experiment. Chronolume reagent (Chrono-log Corporation) was used to monitor ATP release from dense granules.

**P-Selectin surface expression assay**

P-selectin surface expression was measured on platelets from *Vlk*$^{f/f}$ and *Vlk-cKO* mice (n = 3 per genotype) by flow cytometry using FITC–labeled anti-mouse CD62P antibody (BD Pharmingen) in response to indicated concentrations of AYPGKF (Sigma-Aldrich) as previously described.[4] Data is expressed as percentage of P-selectin surface expression compared to stimulated control.

**Hematological analysis of whole blood**

Hematological parameters for 5 mice per genotype were determined using a Hemavet 850FS (Drew Scientific). Parameters analyzed include platelet volume (PLT Vol.); platelet (PLT), white blood cell (WBC) and red blood cell (RBC) count; hemoglobin (Hb) and hematocrit (HCT) levels.

**Tail bleed assay**

Tail bleeding assay was performed as previously described.[5] Briefly, tails from 6-7 mice per genotype were surgically dissected 3 mm from the tip, immersed in buffered saline pre-warmed at 37°C, and time to bleeding cessation was recorded within 20 minutes.

#### Statistics

Statistical significance for all methods, except for laser-induced thrombosis, was determined using Student’s t test. For laser-induced thrombosis, AUC was calculated for platelet and fibrin after laser injury. Mann-Whitney test was used for statistical comparison between genotypes. Statistical analyses were performed with GraphPad Prism 8.

**Supplementary Tables**

**Table S1.** Tyrosine phosphosites mapped to proteins with luminal, secreted, or extracellular domains of transmembrane proteins as annotated in UniprotKB detected in phosphoproteomics of human platelets.

| Protein Name (Gene name) | Phosphosite | Reference(s) |
|---|---|---|
| Fibrinogen alpha chain (*FIBA*) | Y277 | Izquierdo et al., *Thromb Haemost* 2020 |
| Fibrinogen gamma chain (*FIBG*) | Y135 | Izquierdo et al., *Thromb Haemost* 2020 |
| Golgi integral membrane protein 4 (*GOLIM4*) | Y673 | Beck et al., *Blood* 2017 |
| Glycosaminoglycan xylosylkinase (*XYLK/FAM20B*) | Y138 | Izquierdo et al., *Thromb Haemost* 2020 |
| Latent-transforming growth factor beta-binding protein 1 (*LTBP1*) | Y443<br>Y523 | Izquierdo et al., *Thromb Haemost* 2020 |
| Lysosome-associated membrane glycoprotein 1 (*LAMP1*) | Y336 | Izquierdo et al., *Thromb Haemost* 2020 |
| Platelet basic protein (*PPBP*) | Y58 | Izquierdo et al., *Thromb Haemost* 2020; Beck et al., *Blood* 2017 |
| Platelet glycoprotein Ib alpha chain (*GP1BA*) | Y292<br>Y294 | Zahedi et al., *J Proteome Res* 2008 |
| Procollagen-lysine,2-oxoglutarate 5-dioxygenase 2 (*PLOD2*) | Y323 | Zahedi et al., *J Proteome Res* 2008 |
| P-selectin (*SELP*) | Y346 | Izquierdo et al., *Thromb Haemost* 2020 |
| P2X purinoceptor 6 (*P2RX6*) | Y64 | Maguire et al., *Proteomics* 2002 |
| Secreted phosphoprotein 24 (*SPP24*) | Y109 | Izquierdo et al., *Thromb Haemost* 2020 |
| Thrombospondin 1 (*THBS1*) | Y1126 | Izquierdo et al., *Thromb Haemost* 2020 |

**Table S3, related to Figure 3.** Hematological parameters in *Vlk-cKO* mice are comparable to *Vlk*$^{f/f}$ control littermates.

| | ***Vlk***$^{f/f}$ | ***Vlk-cKO*** | **P-value** |
|---|---|---|---|
| PLT Vol. (fL) | 4.2 ± 0.24 | 4.2 ± 0.13 | 0.841 |
| PLT X 10/µL | 692 ± 85.46 | 616 ± 107.2 | 0.250 |
| WBC 10/µL | 4.22 ± 1.23 | 4.58 ± 2.00 | 0.744 |
| RBC $10^6$/µL | 8.64 ± 0.77 | 8.04 ± 0.86 | 0.281 |
| Hb (g/dL) | 12.28 ± 1.05 | 12.90 ± 0.82 | 0.330 |
| HCT (%) | 42.30 ± 3.28 | 38.52 ± 3.75 | 0.128 |

Platelet volume (PLT Vol.); platelet (PLT); white blood cell (WBC) and red blood cell (RBC) count, hemoglobin (Hb) and hematocrit (HCT) levels; data are expressed as mean ± SD; n=5 per genotype.

Supplementary Figures

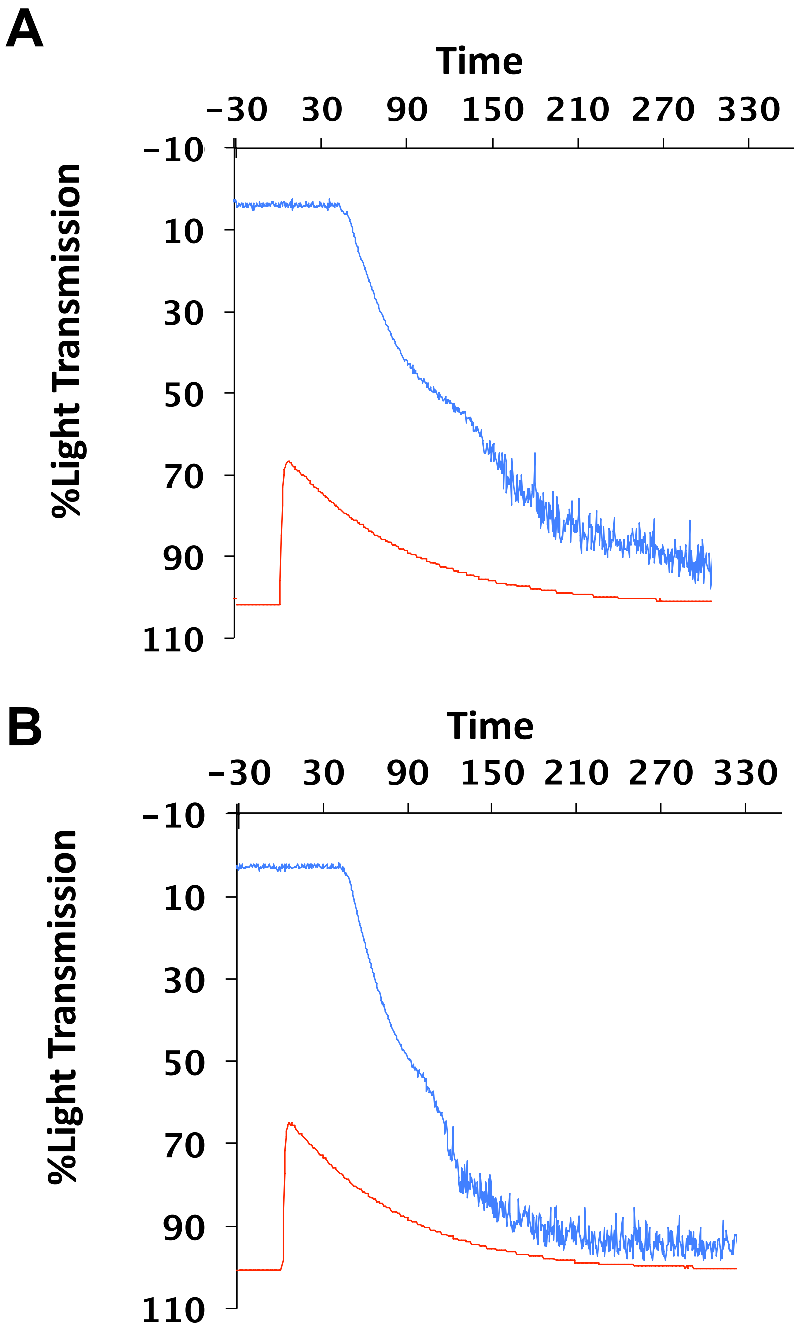

**Figure S1. Platelets from *Pf4*-specific *Vlk-cKO* mice demonstrate normal ADP stores in response to high dose thrombin, related to Figure 3. (A, B)** Representative aggregation (*blue*) and dense granule release (*red*) tracings in response to 5 U/mL thrombin in *Vlk*$^{f/f}$ **(A)** vs. *Vlk-cKO* **(B)** platelets.

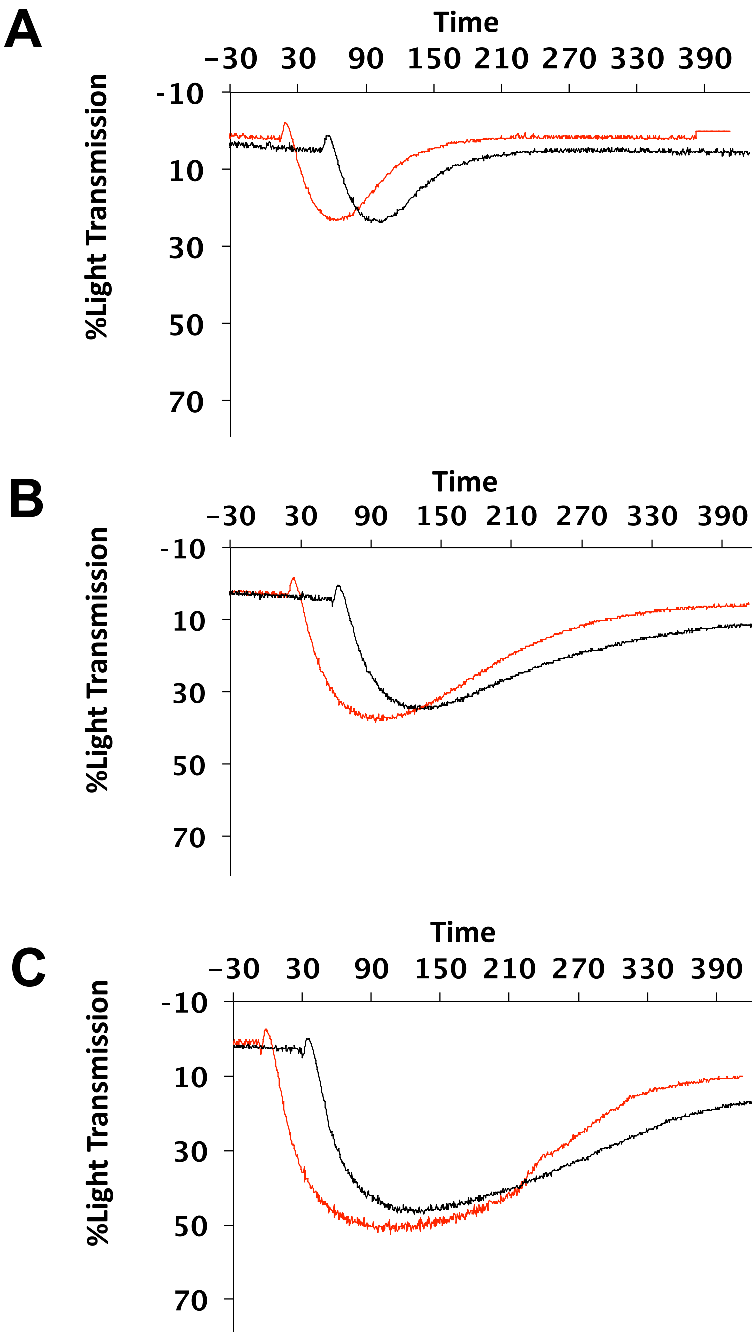

**Figure S2. Platelet aggregation in response to various concentrations of ADP remains unaffected in *Vlk-cKO* platelets compared to control, related to Figure 4. (A-C)** Representative aggregation tracings in response to low (10 μM) **(A)**, intermediate (40 μM) **(B)**, and high (100 μM) **(C)** doses of ADP in *VLK-cKO* platelets (*red*) compared to control (*black*).

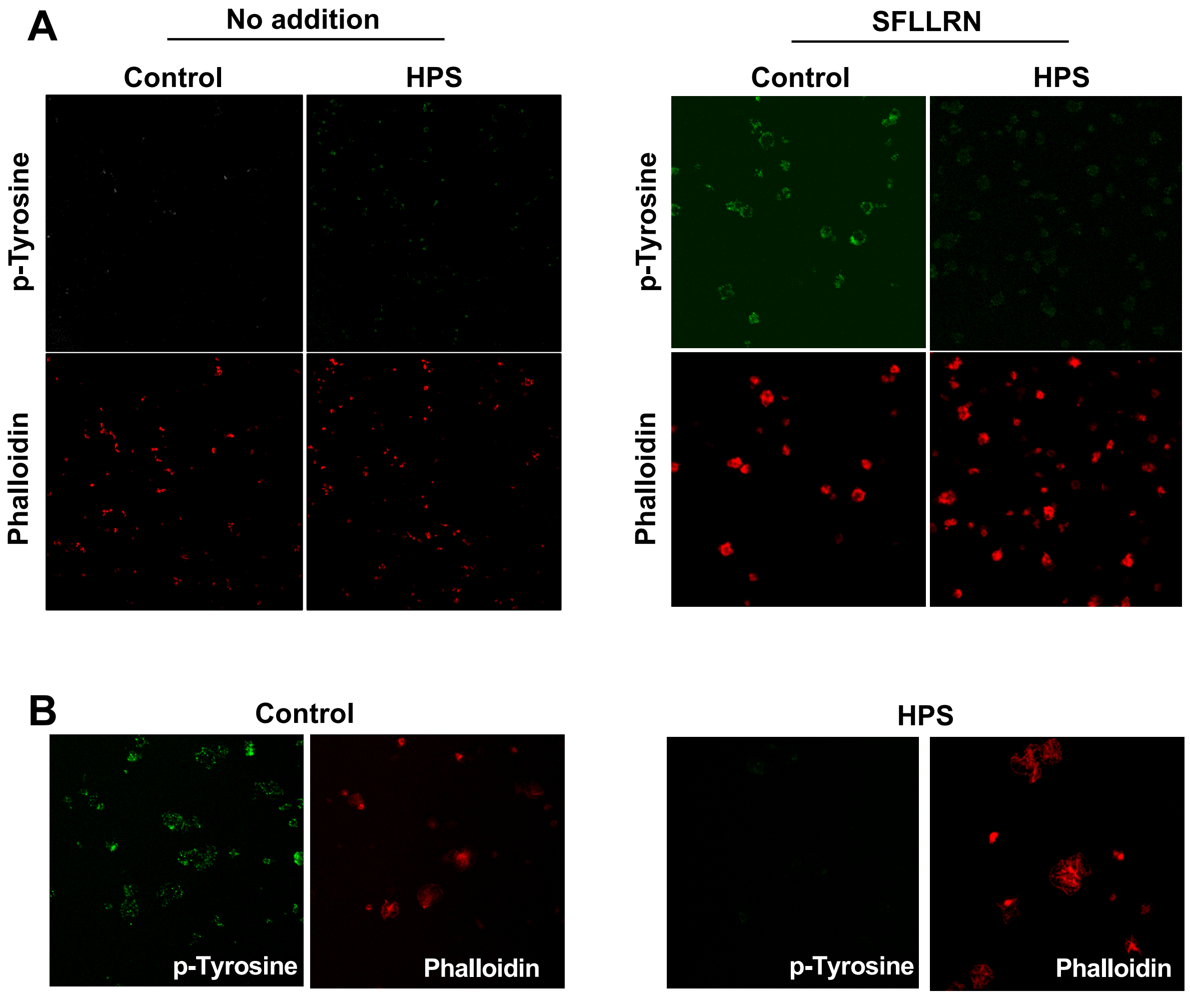

**Figure S3. Impaired extracellular tyrosine phosphorylation in Hermansky-Pudlak syndrome, related to Figure 1. (A)** Platelets from a control subject and a patient with Hermansky-Pudlak syndrome (HPS) were stimulated with 150 μM SFLLRN for 15 minutes. p-Tyrosine on the platelet surface was visualized by staining nonpermeabilized platelets using an anti p-Tyr antibody. Following p-Tyrosine staining, filamentous actin (F-actin) was stained with Alexa Fluor 568 phalloidin diluted in methanol to permeabilize and visualize platelets on slides. (B) Spreading platelets allowed to adhere for 30 minutes on collagen coated coverslips were stained using an anti p-Tyr antibody and Alexa Fluor 568 phalloidin.
